# Caspase-8 is a novel modulator of Homologous Recombination Repair in response to ionizing radiations

**DOI:** 10.1101/2025.05.26.656091

**Authors:** Alessandra Ferri, Claudia Contadini, Claudia Di Girolamo, Claudia Cirotti, Giulia Fiscon, Paola Paci, Marta Marzullo, Maria Pia Gentileschi, Tatsuro Yamamoto, Robert Strauss, Donatella Del Bufalo, Laura Ciapponi, Daniela Barilà

**Affiliations:** Department of Biology, University of Rome "Tor Vergata", 00133 Rome, Italy; Laboratory of Cell Signaling, IRCCS-Fondazione Santa Lucia, 00179 Rome, Italy; Preclinical Models and New Therapeutic Agents Unit, IRCCS Regina Elena National Cancer Institute, 00144 Rome, Italy; PhD Program in Cellular and Molecular Biology, Department of Biology, University of Rome "Tor Vergata", 00133 Rome, Italy; Department of Computer, Control and Management Engineering “Antonio Ruberti”, Sapienza University of Rome, 00185 Rome, Italy; Department of Biology and Biotechnologies "C. Darwin", Sapienza University of Rome, 00185 Rome, Italy; Institute of Molecular Biology and Pathology (IBPM), CNR, 00185 Rome, Italy; Cellular Networks and Molecular Therapeutic Targets Unit, IRCCS Regina Elena National Cancer Institute, 00144 Rome, Italy; Genome Integrity Unit, Danish Cancer Society Research Center, Strandboulevarden 49, Copenhagen, DK-2100, Denmark

## Abstract

Caspase-8 is a cysteine protease historically regarded as anti-neoplastic protein, thanks to its role in apoptosis. However, Caspase-8 expression is retained or even enhanced in several tumors, including glioblastoma (GBM), where it plays pro-tumor functions. We previously reported that it is a negative prognostic factor and contributes to resistance against DNA damaging agents, such as ionizing radiations (IR) and Temozolomide, commonly used in standard GBM treatment. We therefore investigated whether Caspase-8 may sustain DNA repair pathways proficiency in GBM. Here we uncover a novel role of Caspase-8 as promoter of the Homologous Recombination Repair (HRR). Importantly, IR promote Caspase-8 transient nuclear translocation and its recruitment to the chromatin. Moreover, Caspase-8 sustains the expression and the recruitment to the chromatin upon IR of RAD51 and CtIP, two key players of the HRR. Consistently, we identify a synthetically lethal interaction between Caspase-8 and PARP inhibition, that may ameliorate GBM response to IR. Remarkably, by using Caspase-8-/- murine embryo fibroblasts and a Drosophila melanogaster Caspase-8 mutant, we demonstrate that Caspase-8 plays an evolutionary conserved role in DNA repair.

**GRAPHICAL ABSTRACT:** 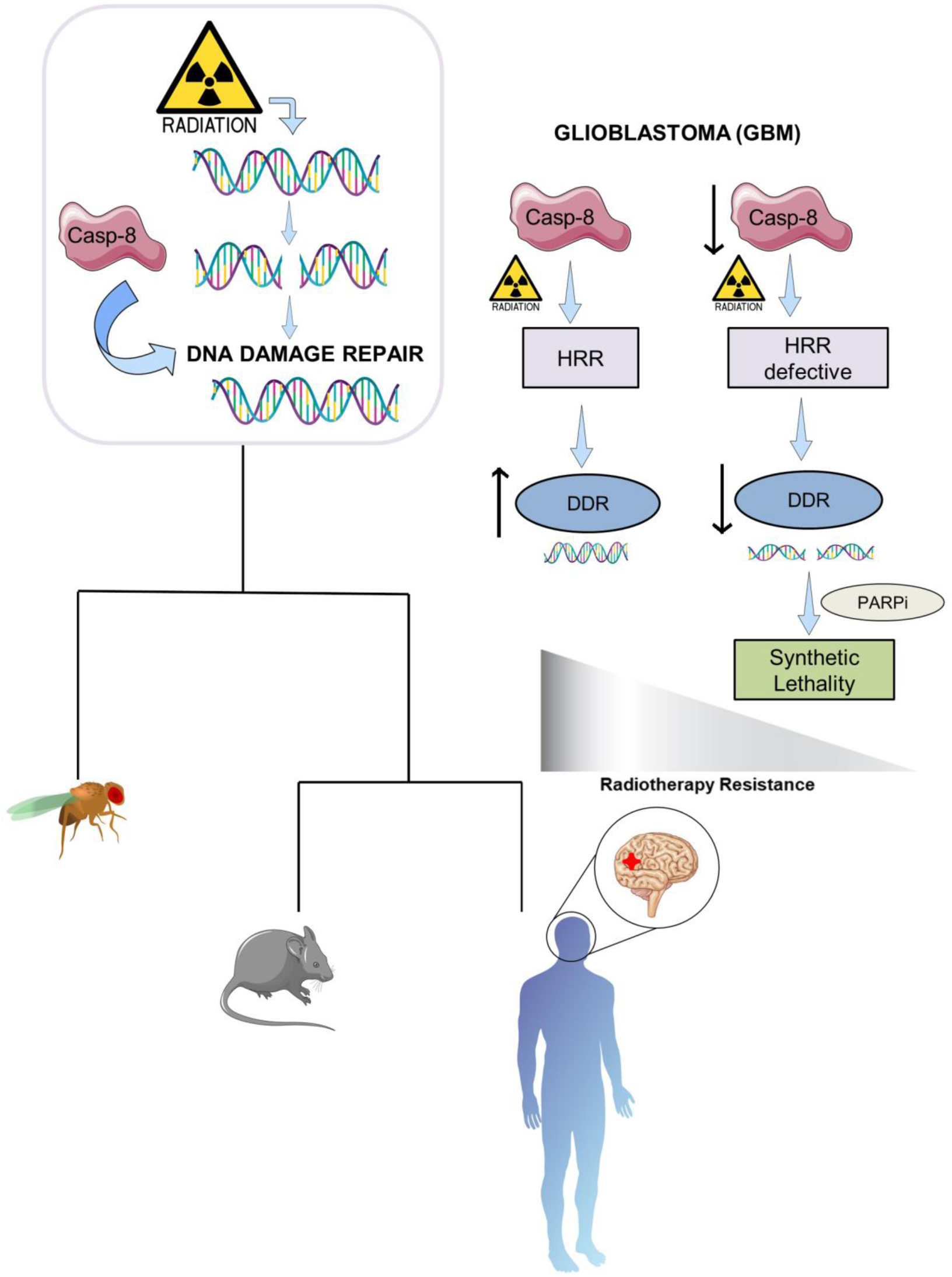

## INTRODUCTION

Glioma is the most frequent brain tumour in adults, accounting for 70% of all the tumours affecting the central nervous system[1]. The most severe form of glioma is glioblastoma (GBM), classified as level IV in the WHO classification. It is extremely deadly, with a dismal prognosis of about one-year survival upon diagnosis[2]. GBM owes its incredible endurance to a multitude of characteristics: its heterogeneity, the abundance of cancer stem cells, its deep infiltrative capacity and overactive DNA repair mechanisms[2].

Over the last fifteen years, no progression has been made at the therapeutic level, with current treatment still consisting mainly of surgical resection followed by cycles of ionizing radiations (IR). The last therapeutic breakthrough saw the introduction of the alkylating drug Temozolomide (TMZ) that improved survival when given in combination with IR[2], but still failed to enhance survival or life quality in a significant matter.

The most severe DNA lesions caused by TMZ and IR are double-strand breaks (DSBs), which are mainly repaired by Homologous Recombination (HR) and Non-Homologous End Joining (NHEJ) pathways. While NHEJ is active throughout the cell cycle and requires minimal homology to ligate the DNA break ends, HR is activated only in S/G2 phases and it is highly accurate as it utilizes the sister chromatid as a template for repair, thereby reducing the risk of misalignment and illegitimate chromosomal rearrangements.[3]. One of the concurrent causes that drive therapy resistance in GBM is the aberrant activation of the DNA repair pathways, which renders DNA damaging treatments ineffective.

Caspase-8 is a cysteine protease associated with the extrinsic pathway of the apoptotic death program and it is therefore considered an unfavorable protein for tumour progression. However, the observation that some tumours, including GBM, unexpectedly upregulate the expression of Caspase-8 suggests that, in specific contexts, Caspase-8 may be beneficial for tumour survival[4,5]. Indeed, non-canonical roles of Caspase-8 have been uncovered over the past fifteen years: its activity in controlling cellular motility and encouraging migration[6], its ability to favour tumorigenesis, anoikis resistance[7] and to promote an inflammatory microenvironment and neoangionesis[8,9]

Furthermore, we previously demonstrated that GBM cells silenced for Caspase-8 expression are more sensitive to TMZ and IR, in line with the inverse correlation found between Caspase-8 expression and the overall survival of GBM patients[8,9]. Based on these observations, we tested whether Caspase-8 expression may reduce GBM cells sensitivity to therapeutic treatments by stimulating the DNA Damage Repair (DDR).

Here we identified in GBM cells Caspase-8 as a prerequisite for DDR, specifically sustaining the expression and the recruitment on chromatin of HR Repair (HRR) factors. We further provide evidence for a nuclear Caspase-8 localization in GBM cellular models both in basal condition and, even more, after IR, demonstrating also its association with chromatin in this context. Remarkably, using murine embryo fibroblasts (MEF) and *Drosophila* model, we show that Caspase-8 sustains DNA repair also in non -tumor and -human contexts and *in vivo,* thereby suggesting that this function is highly conserved.

Overall, we identified an interplay between Caspase-8 and HRR proficiency in GBM cellular models. Since HR-deficient cancer cells are highly sensitive to PARP inhibition[10], we also suggest a synthetic lethality strategy by using PARP inhibitors, such as Olaparib, in combination with IR to enhance the sensitivity of cancer cells defective for Caspase-8 expression to DNA damage induced by IR.

## RESULTS

### Caspase-8 expression sustains DNA damage repair in glioblastoma cellular models

Standard treatment for GBM typically involves DNA-damaging agents, such as IR and TMZ[2]. However, therapy resistance is rapidly acquired through the overactivation of DNA repair pathways [11]. Having previously demonstrated that Caspase-8 expression correlates with reduced sensitivity to IR and TMZ in GBM models[8,9], we tested whether Caspase-8 promotes therapy resistance via modulation of the DDR. To this end, we performed a neutral comet assay to directly measure the amount of fragmented DNA upon treatment with IR. We employed two GBM cell lines (U87-MG and A172), stably expressing either non-targeting shRNA (shCTR) or shRNA against Caspase-8 (shC8) (Fig.1A, S1A). As expected, IR triggers DNA fragmentation independently of Caspase-8 expression, as shown by the length of the comet tails 1h after IR (Fig.1B, S1B). Importantly, 24h post-IR, shC8 cells showed prolonged detection of fragmented DNA, as indicated by the persistence of the comet tail, differently from the respective control cells (Fig.1B, S1B). Moreover, 24h after IR we observed an increased percentage of cells with micronuclei in shC8 compared to shCTR cells (S1C-D). DNA repair upon IR was also assessed through immunofluorescence analysis of pSer139 H2AX (γH2AX), a common marker of DNA damage (Fig.1C, S2A), which triggers the recruitment of several DNA repair proteins to the damage site[12]. Interestingly, although shCTR and shC8 cells exhibit the same amount of γH2AX positive cells 1h after IR, shC8 cells maintain elevated γH2AX foci 24h post-IR. In contrast, shCTR cells are able to repair DNA damage within the same time frame (Fig.1C, S2A). The same result was also obtained using patient-derived GBM stem cells grown as neurospheres (GBM39). These neurospheres were silenced (shC8) or not (shCTR) for Caspase-8 expression (S2B) and their ability to repair DNA damage upon IR was analyzed by immunofluorescence for γH2AX, as above (S2C). Again, Caspase-8 downregulation impairs the repair of damaged DNA, as shown by the persistence of high level of γH2AX still detectable in shC8 cells 24h post-IR treatment (S2C).

**Figure 1.**
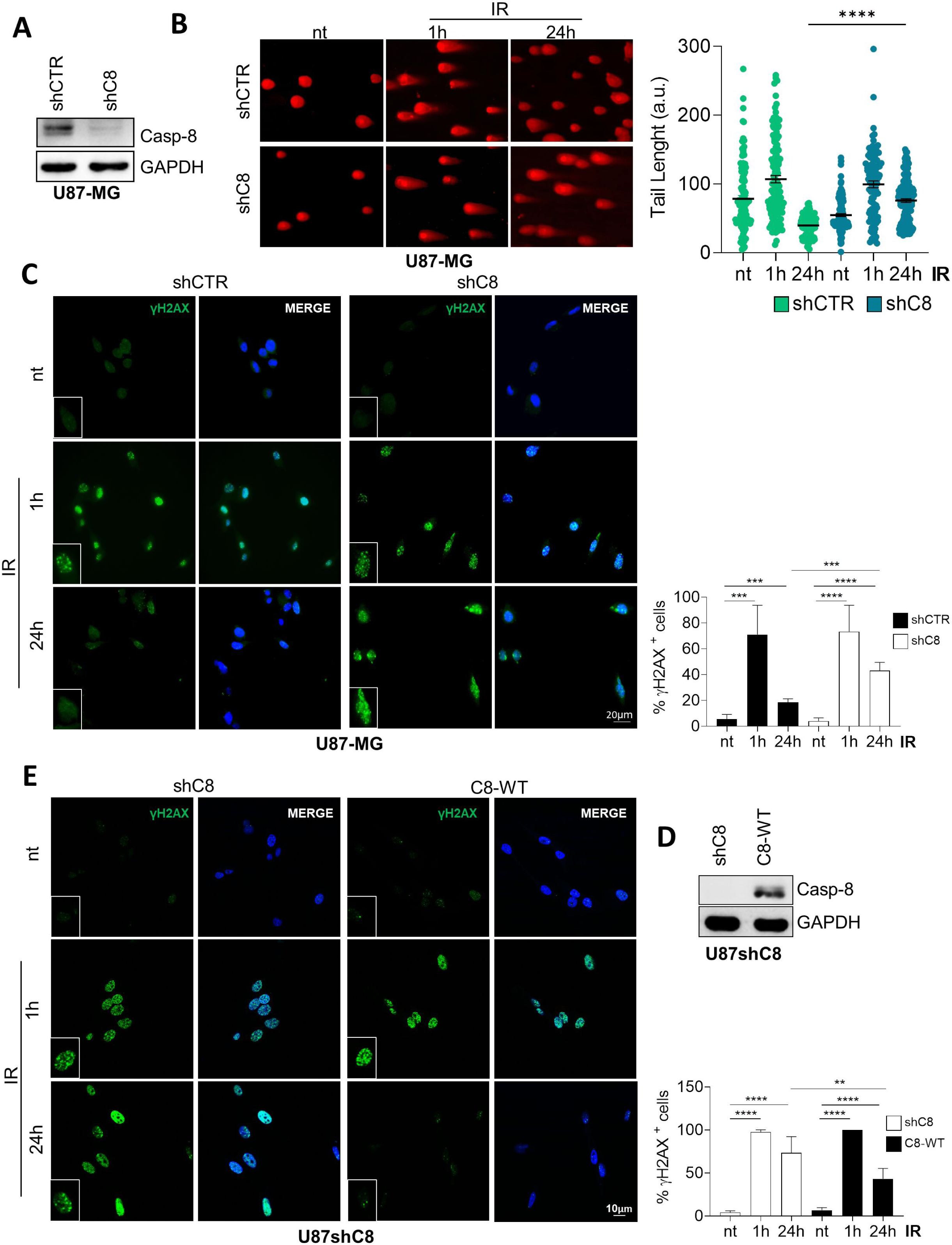
Caspase-8 expression influences DNA repair capacity of GBM cells. **(A)** Immunoblotting on total protein extracts from U87-MG stably silenced for Caspase-8 expression (shC8) or not (shCTR). GAPDH was used as loading control. **(B)** Representative images of Neutral Comet Assay showing Comet tails at time 0 (not treated, nt) and 1h-24h post IR (5Gy) (Left). Graphs representing the median length of the comet (right). **(C)** Immunofluorescence and relative quantification showing the percentage of U87-MG cells (shCTR *vs* shC8) considered positive for the phosphorylation on Ser139 of the histone variant H2AX (n° γH2AX foci>5) at time 0 (not treated, nt) and 1h-24h post IR (5Gy); γH2AX (green), DNA (blue, Hoechst). **(D)** Immunoblotting on total protein extracts from U87-MG stably silenced for Caspase-8 expression (shC8) and stably reconstituted for Caspase-8 expression (C8-WT). GAPDH was used as loading control. **(E)** Immunofluorescence and relative quantification showing the percentage of U87shC8 and U87shC8 cells stably reconstituted with C8-WT considered positive for the phosphorylation on Ser139 of the histone variant H2AX (n° γH2AX foci>5) at time 0 (not treated, nt) and 1h-24h post IR (5Gy); γH2AX (green), DNA (blue, Hoechst). Statistical analysis was performed by the unpaired Student’s t-test (ns= not significant; *P≤0.05; **P≤0.01; ***P≤0.001; **** P≤0.0001). Results represent the mean of three independent experiments ± SD.

We therefore asked whether Caspase-8 expression may impinge on the activation of ATM, the pivotal kinase activated by DNA Double Strand Breaks (DBSs) that triggers the phosphorylation of pSer139 H2AX[12]. Interestingly, ATM phosphorylation levels were not reduced in shC8 cells compared the control ones (S1F-G), which is consistent with the similar amount of damage observed in both shCTR and shC8 cells 1h post-IR.

Remarkably, the stable reconstitution of Caspase-8 expression (U87shC8-C8-WT cells) (Fig.1D), efficiently rescued the DNA damage, as shown by immunofluorescence analysis of γH2AX 1h and 24h post-IR (Fig.1E). Overall, these data support a crucial role for Caspase-8 not in the detection of the damage but rather in the modulation of DDR following the formation of γH2AX foci in response to DNA damage.

### Caspase-8 transiently accumulates in the nucleus and binds chromatin in response to IR

It has been previously reported that Caspase-8 localizes both in the nucleus and cytoplasm in cancer cells[5]. Given the effect of Caspase-8 on DNA repair, we sought to investigate Caspase-8 subcellular localization in our GBM cellular models. Immunofluorescence experiments in U87-MG and A172 cells showed that Caspase-8 localizes both in the nucleus and in the cytoplasm under basal conditions (Fig.2A). Indeed, pretreatment with Leptomycin, a well-known inhibitor of nuclear export, resulted in a strong accumulation of Caspase-8 in the nucleus, further supporting its ability to shuttle between these two cellular compartments (Fig.2B). Interestingly, IR induce a transient accumulation of Caspase-8 in the nucleus, detectable as early as 30 min post-IR, while by 4h post-IR Caspase-8 returned to a diffuse localization, similarly to untreated cells (Fig.2C-D).

**Figure 2.**
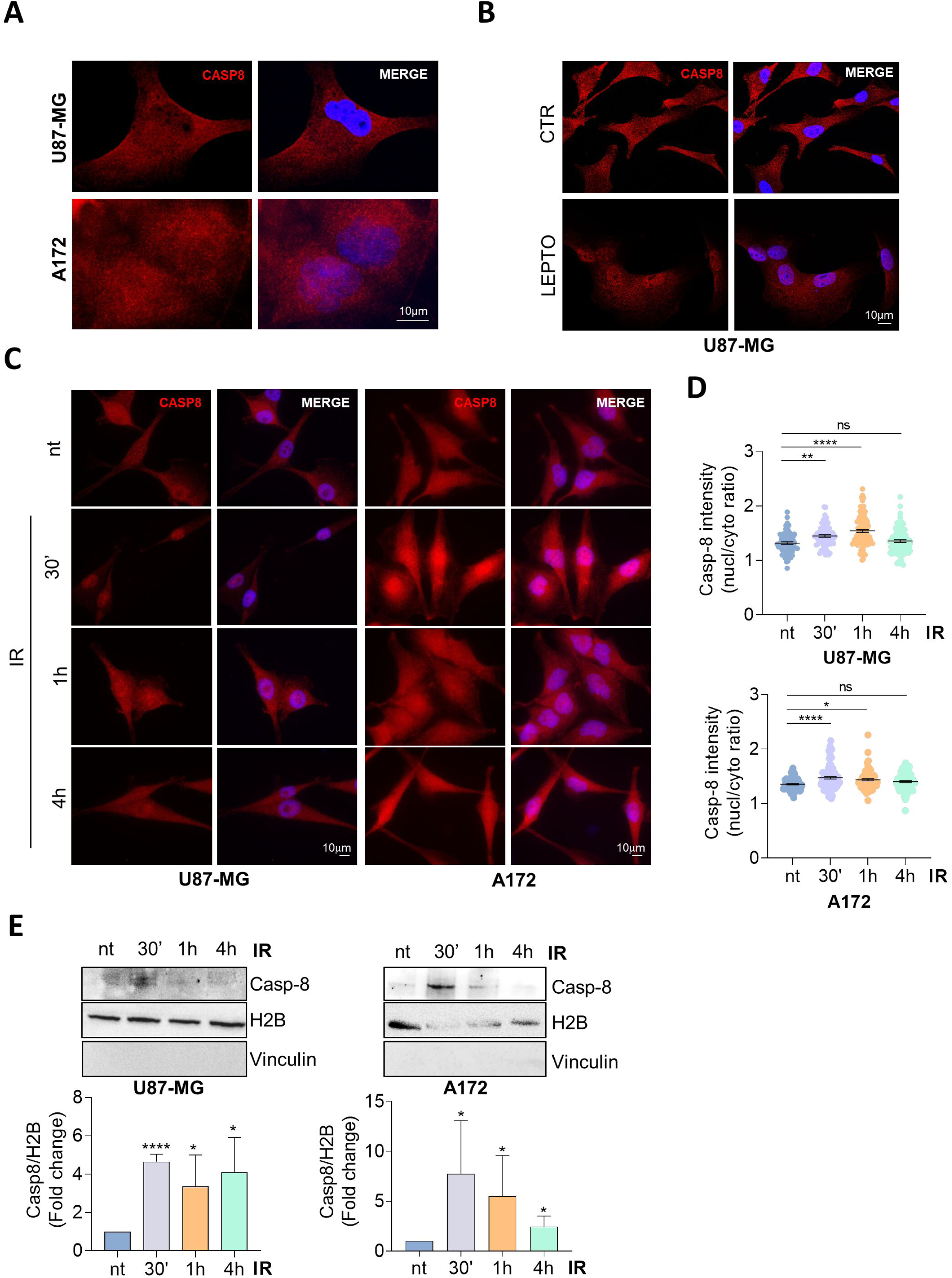
Caspase-8 shuttles between cytosol and nucleus in basal condition and is recruited on chromatin upon IR. **(A)** Immunofluorescence showing a diffuse staining for Caspase-8 (red) in U87-MG and A172 cells. DNA (Hoechst). **(B)** Immunofluorescence showing an accumulation of nuclear Caspase-8 (red) in U87-MG and A172 cells after treatment with Leptomycin B (20 nM) for 3h. DNA (blue, Hoechst). **(C-D)** Immunofluorescence and relative analysis of U87-MG and A172 showing Caspase-8 localization at time 0 (not treated, nt) and 30’-1h-4h post IR treatment (5Gy). Caspase-8 (red), DNA (blue, Hoechst). **(E)** Immunoblotting and relative densitometric analysis of chromatin insoluble portion derived from U87-MG and A172 cells extracted at time 0 (not treated, nt) and 30’-1h-4h post IR (5Gy). H2B was used as chromatin loading control. Statistical analysis was performed by One-way ANOVA statistical test and by the unpaired Student’s t-test (ns= not significant; *P≤0.05; ****P≤0.0001). Results represent the mean of three independent experiments ± SD.

Given its scaffolding functions[13], we also evaluated whether Caspase-8 may be recruited to chromatin in the early phases of DDR. Interestingly, chromatin fractionation experiments revealed that Caspase-8 is recruited to the chromatin 30’ after IR in both U87-MG and A172 cells (Fig.2E). Overall, these data suggest that while in basal conditions Caspase-8 shuttles between the nucleus and the cytosol, upon IR it temporarily accumulates into the nucleus and binds the chromatin, further supporting the idea of a new Caspase-8 function related to DDR.

### Caspase-8 modulates the expression and the chromatin recruitment of RAD51 and CtIP upon IR

To uncover the significance of Caspase-8 expression in GBM cells, we recently performed transcriptomic [8] and proteomic analyses (Data are available via ProteomeXchange with identifier PXD060920) to study the differential gene and protein expression in U87-MG cells genetically silenced (shC8) or not (shCTR) for Caspase-8. The transcriptomic analysis highlighted a significant downregulation of genes involved in inflammation, metabolism and interestingly in IR and DSBs response (S3A). Interestingly, Caspase-8 silencing led to a significant downregulation of genes primarily involved in HRR, but not in NHEJ (Fig.3A), suggesting that Caspase-8 predominantly affects HRR. Consistently, the proteomic analysis revealed a general reduction of DNA Damage response, DDR and DSBs as well (S3B). Analyzing by EnrichR 44 proteins involved in DNA repair’s process, we highlighted a downregulation of factors belonging to HRR (Fig.3B) and involved in DNA binding (Fig.3C). To further confirm the role of Caspase-8 in HRR, we evaluated the basal expression and the recruitment to the chromatin upon IR of CtIP (CtBP-interacting protein) and RAD51 recombinase, two key players of the HRR pathway[14].

**Figure 3.**
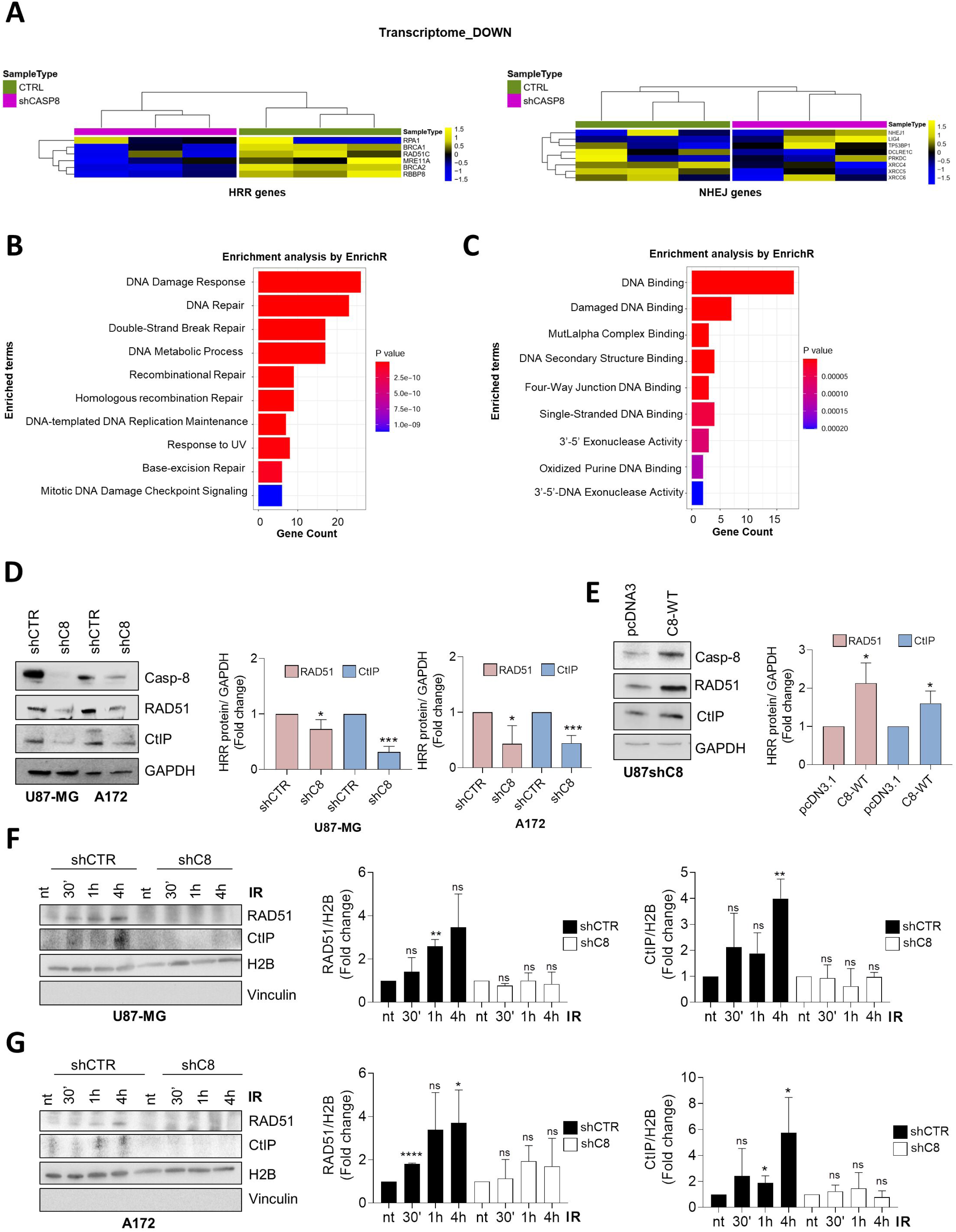
Caspase-8 silencing downregulates DNA Damage repair, Homologous Recombination Repair and impairs chromatin recruitment of RAD51 and CtIP upon DNA damage. **(A)** Heatmaps of expression levels of genes involved in Homologous Recombination Repair (HRR) and Non-homologous end joining (NHEJ) (logarithmic scale) across 6 U87-MG samples grouped by CTRL cells (3 samples, water blue bars) and shCASP8 samples (3 samples, gold bars). A z-score normalization was applied and colors represent different expression levels increasing from blue to yellow. Expression profiles are clustered according to samples (columns) of the data matrix by using Euclidean distance as metrics and complete linkage as clustering algorithm. **(B-C)** Barplot showing the enrichment analysis in Gene Ontology Biological Processes (B) and Molecular Function (C) for the 44 genes involved in DNA repair’s process. Bars length refers to the number of gens annotated in each category, whereas bars color refers to the p-value of the functional enrichment analysis performed by querying EnrichR tool. **(D)** Immunoblotting and relative densitometric analysis showing CtIP and RAD51 protein level in shCTR and shC8 cells of U87-MG and A172 cell lines. GAPDH was used as loading control. **(E)** Immunoblotting and relative densitometric analysis showing CtIP and RAD51 protein level in U87-MG shC8 and shC8 cells transiently reconstituted for Caspase-8 expression. GAPDH was used as loading control. **(F-G)** Immunoblotting and relative densitometric analysis of chromatin insoluble portion derived from shCTR and shC8 (U87-MG and A172) at time 0 (not treated, nt) and 30’-1h-4h post IR treatment (5Gy). H2B was used as chromatin loading control. Results represent the mean of three independent experiments ± SD. Statistical analysis was performed by the unpaired Student’s t-test (ns= not significant; *P≤0.05; **P≤0.01; ***P≤0.001; ****P≤0.0001).

According to the transcriptomic analyses, shC8 cells showed a significant reduction of both CtIP and RAD51 expression compared to shCTR (Fig.3D). Remarkably, the reconstitution of Caspase-8 expression (shC8-C8-WT) rescued CtIP and RAD51 expression levels (Fig.3E). More interestingly, Caspase-8 silencing impaired their recruitment to the chromatin upon IR (Fig.3F-G), thus suggesting that Caspase-8 sustains their expression in basal condition as well as their activity during DDR.

Overall, these data highlight a role of Caspase-8 in promoting the expression and the chromatin recruitment upon DNA damage of factors involved in HRR, therefore enhancing DDR proficiency.

### Caspase-8 sustains the Homologous Recombination Repair pathway therefore promoting DNA damage resolution

To further verify whether Caspase-8 impinges on HRR, we performed immunofluorescence analysis of shCTR and shC8 cells upon IR to detect BRCA1 foci formation, a well-established marker of HRR activation involved in CtIP activation and RAD51 regulation in response to DNA damage[3,15]. HRR is an error-free repair pathway that occurs in the S/G2 phases of the cell cycle as it requires the sister-chromatid to repair DSBs. We used Cyclin A as marker of the S/G2 phases to selectively count BRCA1 foci only in Cyclin A^+^ cells (Fig.4A,C, S4A,C). As expected, a significant increase of BRCA1 foci number in shCTR cells was detected 1h after DNA damage induction, followed by its reduction 24h later, consistently with the efficient activation of the HRR and the subsequent resolution of the damage. On the other hand, shC8 cells displayed an overall dampened recruitment of BRCA1 on DNA, consistently with impaired activation of the HRR (Fig.4A,C, S4A,C). We then asked whether to overcome the defect of HRR, shC8 cells may trigger a stronger activation of the NHEJ. To this aim, immunofluorescence experiments were performed to detect 53BP1 as a marker for NHEJ activation[16]. Importantly, Cyclin A was used to mark cells in the S/G2 phases of the cell cycle, when the HRR and the NHEJ are competing, and 53BP1 foci were counted only in Cyclin A^+^ cells. As before, DNA damage was evaluated 1h and 24h post-IR. Interestingly, no significant difference in the number of 53BP1 foci was detected between shCTR and shC8 1h after IR, prompting us to conclude that shC8 cells do not over-activate the NHEJ to compensate for the reduced HRR (Fig.4B,D, S4B,D). Consistently, independently of Cyclin A positivity, both shCTR and shC8 cells recruit 53BP1 in a comparable manner, thus supporting the conclusion that Caspase-8 expression does not significantly modulate NHEJ (S4E,F).

**Figure 4.**
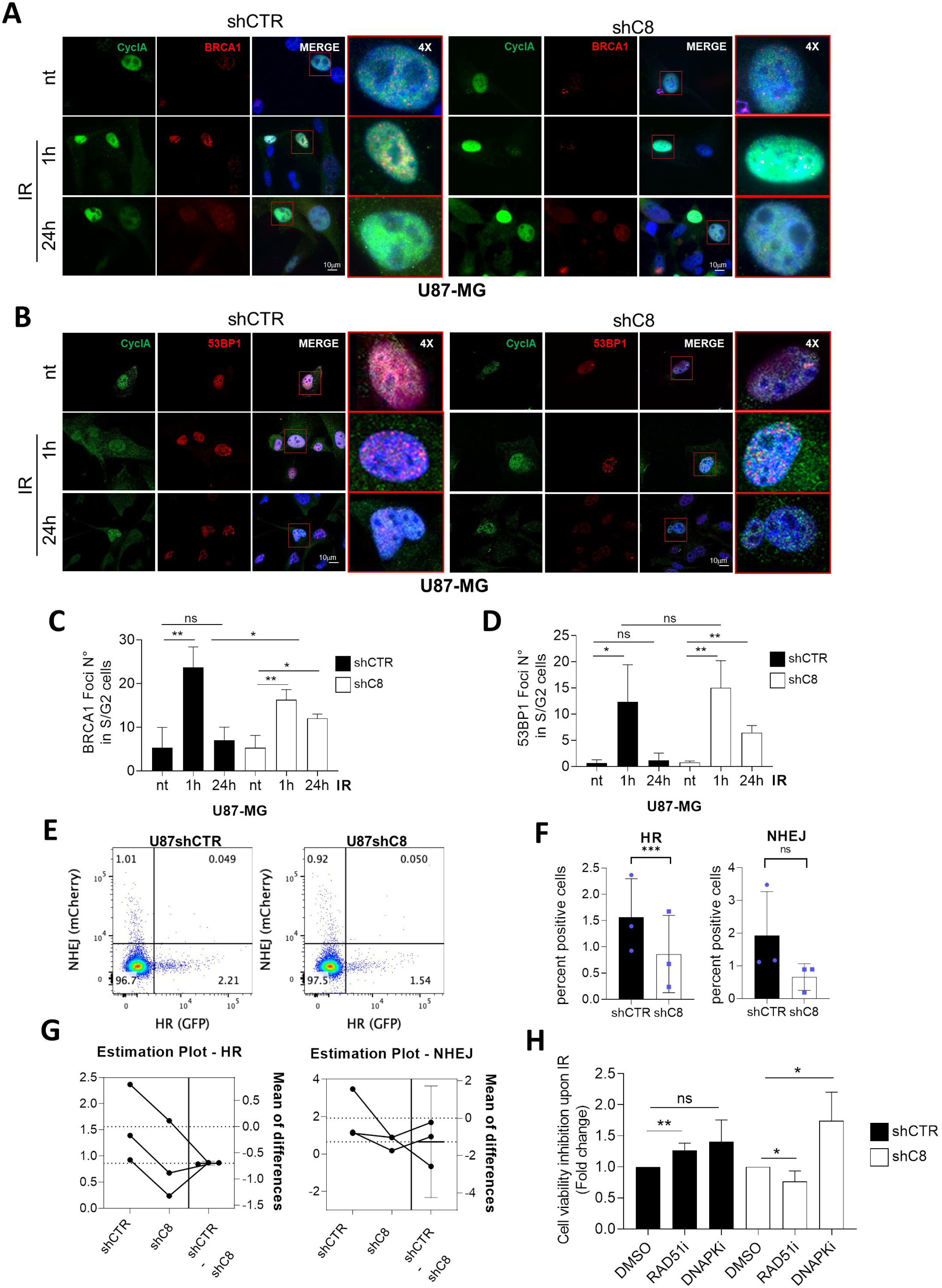
Caspase-8 exerts its effect on DNA repair influencing proficiency of the Homologous Recombination Repair pathway. **(A-B)** Immunofluorescence analysis performed on U87-MG shCTR and shC8 cell lines to detect Cyclin A (green) and BRCA1 (**A,** red) or 53BP1 (**B**, red) foci at time 0 (not treated, nt) and 1h-24h post IR (5Gy). DNA (blue, Hoechst). **(C-D)** Bar-graph indicating the average BRCA1 or 53BP1 foci number in CycA^+^ cells. **(E)** Representative plots of traffic light reporter assay in U87shCTR and U87shC8 cells 72 hours after induction. Homologous Recombination (HR) was assessed by reconstitution of GFP, while non-homologous end joining (NHEJ) was visualized via mCherry. **(F)** Mean percentage GFP- (HR) and mCherry- (NHEJ) positive cells with standard deviation of three independent experiments. The correlation between independent experiments is visualized by estimation plots **(G)**. **(H)** MTS assay measuring U87-MG cell viability after 72h from IR. U87shCTR and shC8 cells were pre-treated 1h before IR with the indicated inhibitors (RAD51i, 27μM; DNAPKi, 2μM) and then irradiated or not with 10Gy IR. Statistical analysis was performed by the unpaired Student’s t-test (ns= not significant; **P≤0.01; ***P≤0.001). Results represent the mean of three independent experiments ± SD.

Furthermore, we assessed the DNA repair pathway choice by using the traffic light reporter system [17] in U87shCTR and shC8 cells. Traffic light reporter assay provides a flow cytometric readout of HRR (GFP) or NHEJ (mCherry) upon DNA damage. In accordance with our previous findings, Caspase-8 silencing consistently reduced HRR capacity, while NHEJ was only affected in a random manner (Fig.4E-G).

Finally, U87-MG cells depleted or not for Caspase-8 expression were irradiated in the presence of RAD51 (RAD51i) or DNAPK (DNAPKi) inhibitors, to selectively block the HRR and the NHEJ, respectively. As expected, both inhibitors slightly increased the sensitivity of shCTR cells to IR (Fig.4H). Remarkably, U87shC8 cells exhibited greater sensitivity to co-treatment with the NHEJ inhibitor rather than with HR inhibitor (Fig.4H), consistent with our earlier observation that U87shC8 cells have impaired HRR and therefore depend more heavily on NHEJ for DNA repair. Overall, these data strongly confirm a role for Caspase-8 in the modulation of HRR functionality.

### Caspase-8 expression sustains DRR in non-transformed *in vitro* and *in vivo* models

We further investigated this novel Caspase-8 function in a non-tumoral context, by using murine embryonic fibroblasts (MEF), eighter *wild-type* (WT) or Caspase-8 *knock-out* (C8 -/-). As before, we analyzed the phosphorylation of γH2AX by performing immunoblotting experiments in non-irradiated condition and 1h-24h after IR (5Gy). As shown in Fig.5A, 24h after IR γH2AX levels were higher in C8 -/- MEFs compared to WT, suggesting that the absence of Caspase-8 impairs DDR functionality. This data was confirmed by immunofluorescence analyses of the percentage of γH2AX positive cells following irradiation (Fig.5B). Indeed, 24h after irradiation. the proportion of γH2AX positivecells was significantly higher in C8 -/- MEFs compared to WT.

**Figure 5.**
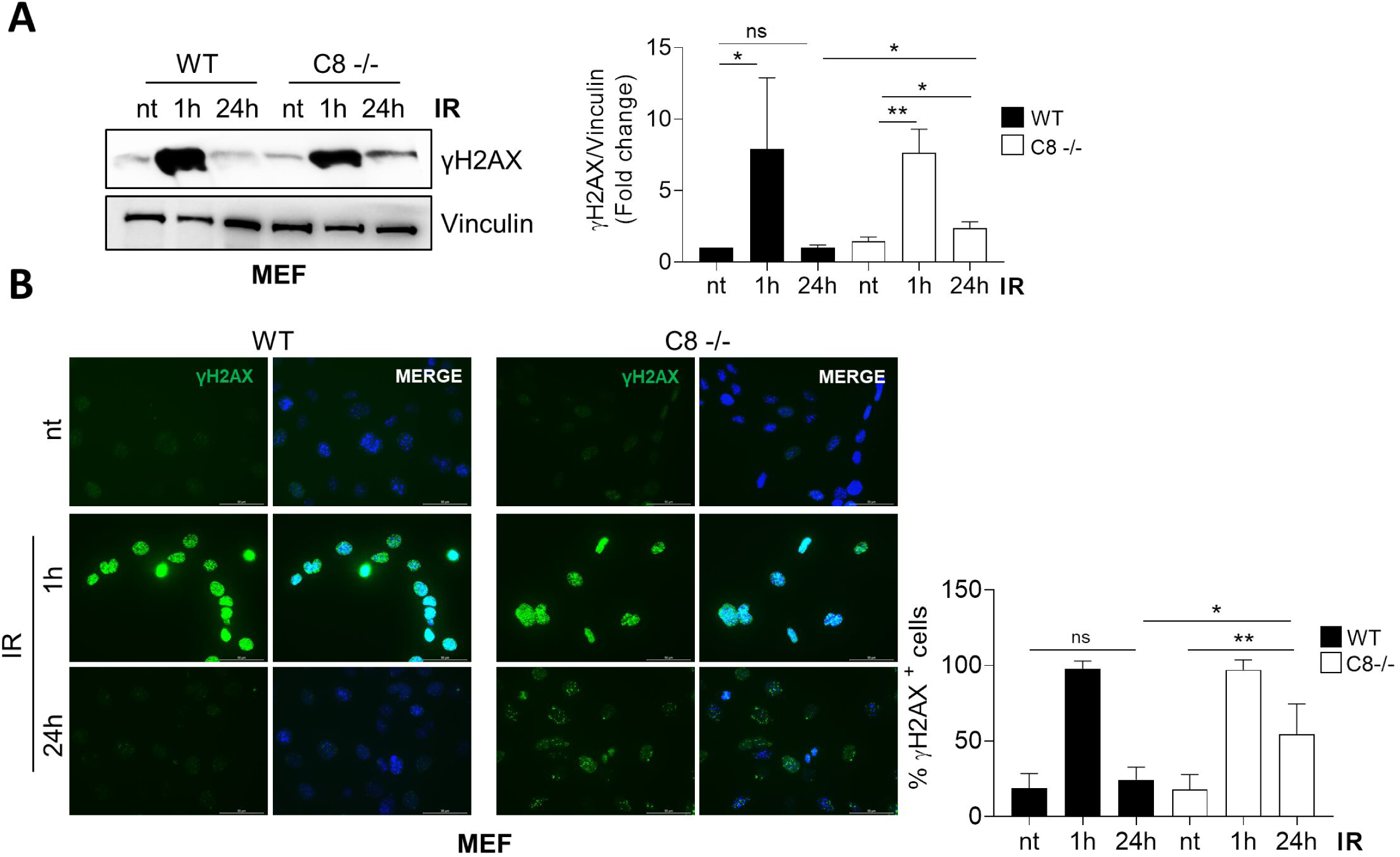
Caspase-8 expression sustains DNA damage repair in murine embryo fibroblasts cellular model. **(A)** Immunoblotting and relative densitometric analysis showing γH2AX levels on total protein extracts from wild-type (*WT*) or Casp-8 knock-out (C8 -/-) MEFs. Vinculin was used as loading control. **(B)** Immunofluorescence and relative quantification showing the percentage of cells considered positive for the phosphorylation on Ser139 of the histone variant H2AX (n° γH2AX foci>5) at time 0 (not treated, nt) and 1h-24h post IR (5Gy); γH2AX (green), DNA (blue, Hoechst). Statistical analysis was performed by the unpaired Student’s t-test (ns= not significant; *P≤0.05; **P≤0.01). Results represent the mean of three independent experiments ± SD.

These data suggest that this newly identified non canonical function of Caspase-8 in DDR is also present in a non-tumoral context and isplausibly conserved through evolution. To verify this evolutionary aspect, we conducted the same experiments using a *Drosophila melanogaster* Caspase-8 mutant as model system. The *Drosophila* Caspase-8 orthologue, Dredd, has been shown to have a conserved role in apoptosis[18] and it is also integral to inflammatory signalling activated by bacterial infections, acting as a driver of NF-κB-dependent immune responses[19,20]. Specifically, *D44* mutants carry mutations in the DED domain, rendering them Caspase-8 defective mutants. These mutations significantly impair Dredd ability to interact with NF-κB and trigger the immune response[20].

Taking advantage of this mutant, we evaluated the ability of *Drosophila* Dredd to mediate DNA repair. *Drosophila* possesses a single histone H2A variant, H2Av, which is thought to fulfill the roles of both mammalian H2A.Z and H2A.X in transcriptional regulation and DNA damage response[21]. Thus, to analyze the kinetics of X-ray-induced phosphorylation of H2AvD at S137 (γH2Av), we irradiated *dredd^Δ44/Δ44^* mutants and *wild-type Drosophila* larvae with X-rays at 5Gy. The levels of γH2Av in larval brains were assessed at various post-irradiation time points (post-IR, 10’, 30’, 1h and 2h) using both immunoblotting and immunofluorescence techniques.

Immunoblotting analysis revealed that irradiated *wild-type* brain extracts display high levels of γ- H2Av at 10- and 30-minutes post-IR but decreased at 1- and 2-hours post-IR, consistent with canonical DDR kinetics[22]. In contrast, in *dredd* mutants brain extracts, high levels of γ-H2Av were retained at 1- and 2-hours post-IR, suggesting that the defective Dredd impairs DDR process (Fig.6A). DNA repair kinetics following 5Gy irradiation were assessed in larval brains also through immunofluorescence analysis, quantifying the number of γ-H2Av foci per cell (Fig.6B-C). Consistent with the immunoblotting data, both *wild-type* and *dredd^Δ44/Δ44^* mutant brains showed similar numbers of γ-H2Av-positive cells at 10- and 30-minutes post-irradiation. However, at 1- and 2-hours post-irradiation, *dredd ^Δ44/Δ44^* cells retained high levels of γ-H2Av foci, in contrast to *wild-type* cells, which effectively repaired the DNA damage. This difference was most pronounced 2h post-irradiation, with *wild-type* cells showing an average of 6.6 foci per cell compared to 8.7 foci per cell in *dredd* mutant cells. These data emphasize the crucial role of the Drosophila Caspase-8 orthologue Dredd in DDR signalling and highlight the *in vivo* conservation of this non-canonical function.

**Figure 6.**
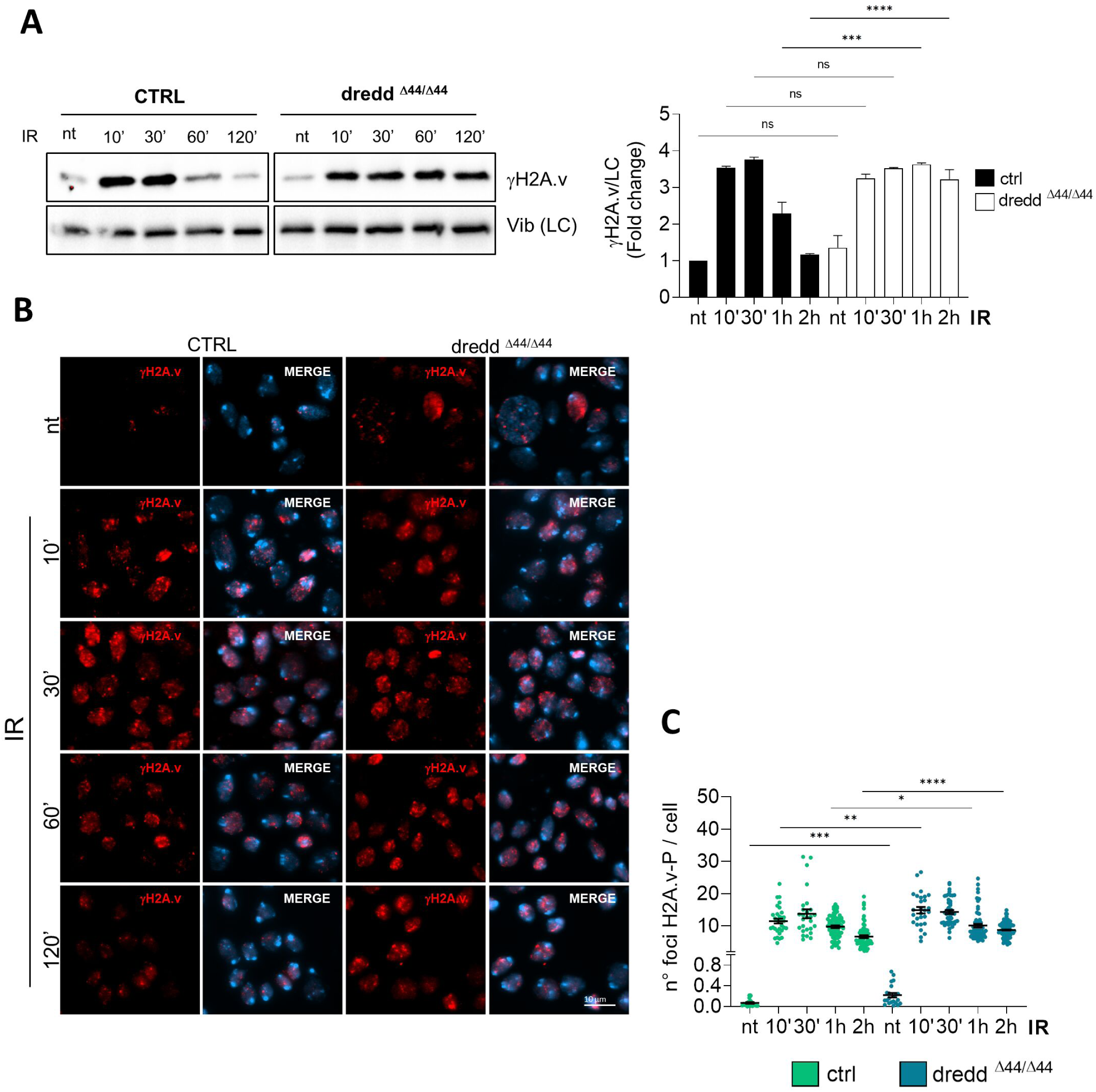
Caspase-8 expression sustains DNA damage repair in Drosophila melanogaster. **(A)** Immunoblotting and relative densitometric analysis showing H2A.v phosphorylation (γH2A.v) levels on total protein extracts from *dredd^Δ44^*mutants (*dredd^Δ44/Δ44^*) or wild-type (*WT*) *Drosophila* larval brains. Vibrator was used as loading control (LC). **(B-C)** Immunofluorescence and relative quantification showing the number of γH2A.v foci in nuclei from *dredd^Δ4z4^* mutants or *WT* larval brains at time 0 (not treated, nt), or 10’, 30’, 1h and 2h post IR, (5Gy); γH2A.v (red), DNA (DAPI, blue). Statistical analysis was performed by the non-parametric Kruskal-Wallis test (ns= not significant; *P≤0.05; **P≤0.01; ***P≤0.001; ****P≤0.0001). Each dot represents a specific field. Number of analyzed brains ≥ 3.

### Caspase-8 silencing in GBM cells confers sensitivity to PARP inhibition

We have previously demonstrated that GBM cells expressing Caspase-8 are more resistant to IR compared to the ones silenced for its expression[8].

According to the role of Caspase-8 in HRR and to the identification of the synthetic lethal interaction between PARP inhibitors and HR defects, we speculated that Caspase-8 expression may represent a novel vulnerability to sensitize irradiated cells to Olaparib, a competitive PARP-1/2 inhibitor. To deepen this issue, we compared GBM shCTR and shC8 cell viability 72h post the following treatments: DMSO (nt), Olaparib (10 μM), IR (10Gy), and Olaparib combined with IR. In line with our hypothesis, shC8 cells are more affected then shCTR cells by the co-treatment Olaparib plus IR, as shown by MTS assay (Fig.7A-B) and cytofluorometric analysis of cell death (Fig.7C-D). Overall, Caspase-8 may represent a novel DDR factor whose mutation or loss of expression confers sensitivity to PARP inhibitors in combination with radiotherapy[23].

**Figure 7.**
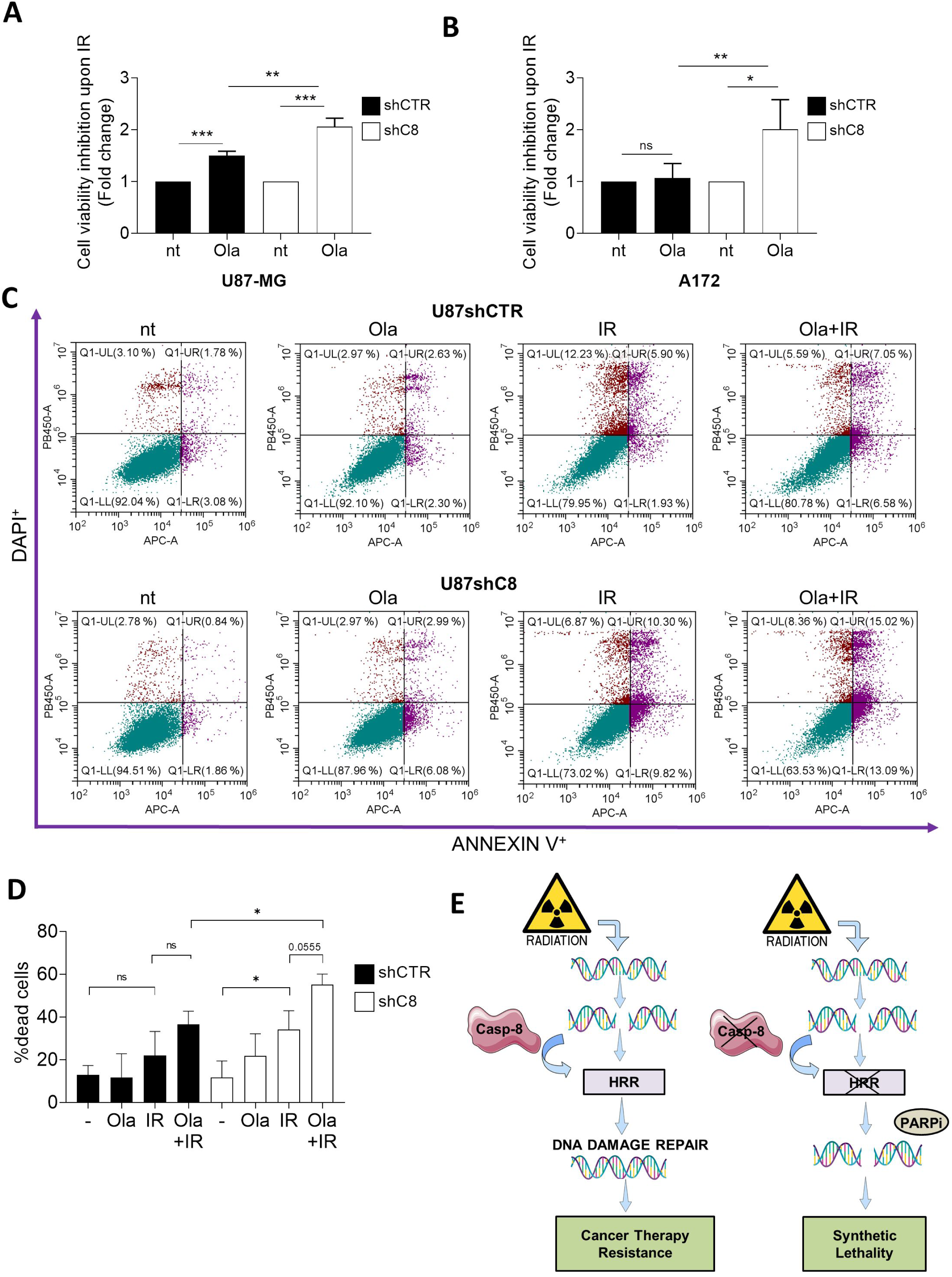
Caspase-8 silencing promotes synthetic lethality in GBM cellular models. **(A-B)** MTS assay measuring cell viability in shCTR and shC8 cell (derived from U87-MG and A172) after 72h from IR (10Gy). 2h before irradiation shCTR and shC8 cells were pre-treated with Olaparib (10μM). Results represent the mean of two (U87-MG) and three (A172) independent experiments ± SD. **(C-D)** Representative flow cytometry dot plot graphs and relative histogram of the percentage of dead cells upon AnnexinV-DAPI staining of U87shCTR and shC8 cells after 72 h irradiation (IR 10 Gy) pre-treated 2h before IR with Olaparib (10μM). Results represent the mean of three independent experiments ± SD. Statistical analysis was performed by One-way ANOVA statistical test for each cell lines compared to the untreated condition. Unpaired t-test was performed for shC8 Ola+IR condition compared to the respective shCTR condition. (ns= not significant;* p<0.05; **p<0.01; ***p<0.001; ****p<0.0001). **(E)** Schematic illustration of Caspase-8 function in DNA damage Repair in glioblastoma. Left panel: Caspase-8 expression in cancer supports Homologous Recombination Repair (HRR) and therapy resistance. Right panel: Loss of Caspase-8 expression results in defective HRR and enhances cancer cells sensitivity to combined treatments with IR and PARP inhibitors.

## DISCUSSION

GBM is the most aggressive primary brain tumor in adults, characterized by substantial morbidity and mortality, which make it a devastating disease. The standard of treatment for GBMs is surgery, followed by chemo-therapy with TMZ and radio-therapy, which share common pathways to cell death, inducing DNA damage either directly or indirectly[2]. Unfortunately, the highly infiltrative nature of GBM allows the tumour to extensively invade the surrounding brain parenchyma, making complete surgical resection unfeasible[24], while the hyper-activation of the oxidative stress response and the DNA damage repair system allow cancer cells to overcome the DNA damage induced by IR and TMZ[24,25]. Moreover, GBM is a very heterogeneous tumour and indeed, based on the gene expression profile, four different sub-types (pro-neural, neural, classic and mesenchymal), each characterized by specific mutations and different susceptibility to therapies have been identified[26]. Among these, the mesenchymal subtype has been identified as the most aggressive and therapy resistant[27,28]. In addition, treatment with IR and TMZ stimulates pro-neural to mesenchymal transition[29], selecting tumor cells with higher IR resistance, thus resulting in a poorer prognosis[26,30,31]. Given the high intra- and inter-tumor heterogeneity, the identification of new molecular features or signatures to stratify patients, together with innovative combinatory strategies, is necessary to fight this aggressive tumour and improve patients outcome. Interestingly, the mesenchymal sub-type, typically characterized by PTEN and NF1 loss, is also characterized by high levels of Caspase-8 expression[26].

Caspase-8 is a cysteine protease usually known for its role in the extrinsic apoptotic pathway and coherently, its expression levels are commonly reduced in many cancer types as a mechanism to escape apoptotic cell death. Therefore, the elevated expression of Caspase-8 in the mesenchymal subtype was unexpected and prompted investigations into how Caspase-8 might contribute to tumor survival.

The discovery of non-canonical roles identified Caspase-8 as a central regulator of cell motility[6], tumorigenesis[7], inflammatory tumour microenvironment (TME) and neo-vascularization[8,9,32]. Interestingly, we have previously uncovered an inverse correlation between high Caspase-8 levels and GBM patients’s survival[9]. Moreover, we have demonstrated that GBM cells silenced for Caspase-8 expression display increased sensitivity to TMZ[9] and IR[8].

In this study, we investigated the mechanism underlying the connection between Caspase-8 and therapy resistance, with a focus on the DNA damage response (DDR) system. Notably, we uncovered an unexpected and significant role for Caspase-8 in promoting the DNA repair capacity of GBM cells, particularly influencing the HRR. Interestingly, Boege et al. previously reported the involvement of Caspase-8 in the DDR by affecting the initial formation of γH2AX foci in mice hepatocytes[33]. Here, we demonstrate that Caspase-8 expression does not influence the initial response to DNA damage but instead sustains the expression of key DDR factors, primarily those involved in HRR, and facilitates their recruitment to sites of DNA damage, thereby promoting efficient repair. Indeed, genetic inhibition of Caspase-8 expression leads to a dramatic downregulation of RAD51 and CtIP expression and more importantly severely impinges on their recruitment to the chromatin in response to IR treatment. We found that Caspase-8 is capable of shuttling between the cytosol and the nucleus, and, notably, following IR, we observed a transient accumulation of Caspase-8 within the nuclear compartment where it can associate to the chromatin. Interestingly, this subcellular relocalization of Caspase-8 upon DNA damage, has been previously reported[5]. In melanoma cells, DNA damaging agents promote Caspase-8 accumulation into the nucleus where it cleaves the de-ubiquitinase USP28 (Ubiquitin Specific Peptidase 28), therefore promoting p53 ubiquitination and proteasomal degradation[5].

Future experiments will clarify both the molecular mechanisms that modulate Caspase-8 shuttling as well as its ability to modulate DNA repair both in GBM cells as well as in other tumors. Interestingly, we provide evidence for a conserved role of Caspase-8 as promoter of DNA repair also in non - tumorigenic contexts *in vitro* and *in vivo.* Using the *Drosophila Melanogaster* model system, in which Caspase-8 plays apoptotic and non-apoptotic functions[18-20], we demonstrate that mutant larvae defective for Caspase-8 expression, display a significant accumulation of DNA damage upon IR compared to the *wt* counterpart, further supporting a key role of Caspase-8 as novel DNA repair player. Furthermore, using Caspase-8 -/- mouse embryo fibroblasts, we provide evidence for a role of Caspase-8 as modulator of DNA repair also in a non-tumoral context, where it has been previously demonstrated its involvement in the maintenance of chromosome stability[34]. In this context Caspase-8 together with RIPK1 has been suggested to modulate PLK1 activity therefore surveilling chromosome segregation. These results provide evidence for a role of Caspase-8 expression as safe-guardian of genomic instability[34]. Our data, highlight for the first time a direct role of Caspase-8 in the modulation of DNA repair, that may further support genome stability. Importantly, we can speculate that this function may have a dual role in cancer. Indeed, on one side Caspase-8 expression may counteract the accumulation of DNA damage and therefore protect cells from neoplastic transformation, but on the other side Caspase-8 by sustaining DNA repair may trigger cancer cell resistance to radio and chemotherapy.

In this regard, the identification of Caspase-8 as a promoter of HRR pointed to Caspase-8 expression levels as a novel marker to select those tumors that may be more sensitive to PARP inhibitors. Indeed, we could show that genetic inhibition of Caspase-8 expression, which results in defective HRR significantly enhances GBM cells sensitivity to the combined treatment with IR and Olaparib (Fig.7E). We therefore suggest that the evaluation of Caspase-8 expression levels may represent ameliorate patient stratification and improve the selection of the best therapeutic strategy.

## Materials and Methods

### Cell Culture

Glioblastoma cell lines (U87-MG, A172) and murine embryo fibroblasts (MEFs) were grown in Dulbecco Modified Eagle Medium (DMEM) (Sigma Aldrich), supplemented with 10% south-american foetal bovine serum (Sigma Aldrich), 1% L-Glutamine (Sigma Aldrich), streptomycin 10mg/ml, penicillin 1000U/ml (Sigma Aldrich). Cells were cultured at 37°C and 5% CO_2_. Glioblastoma patient-derived GBM39 cells were cultured as neurospheres in suspension in F-12 Medium supplemented with B27 Supplement (Gibco 17504044, 50x), EGF (20ng/ml), and βFGF (10ng/ml) as previously described[8]. All GBM cells were routinely tested negative for mycoplasma contamination.

### Stable Caspase-8 knock-down in GBM cell lines

Stable interference of Caspase-8 was obtained through expression of two short-hairpin RNA (shRNA) as described in[8].

### Stable or transient reconstitution of Caspase-8 *wild-type*

For the stable reconstitution of Caspase-8-WT, shC8 cell lines (obtained using E42 clone, see below) were infected with lentiviral pLV-EF1a IRIS vector encoding for Caspase-8-WT carrying the mutations that do alter amino-acid sequence but also confer resistance to the above interference: 5’-GCacTtATGcTtTTCCAacGt-3’.

For the transient reconstitution of Caspase-8-WT, we transfected shC8 cells with the empty vector pcDNA3.1 and pcDNA3.1 Caspase-8 by using Lipofectamine2000 (Invitrogen) according to the manufacture instructions.

### Antibodies and other reagents

Anti-BRCA1 (Santa Cruz Biotechnology, 1:250 sc-6954), anti-Caspase-8 (MBL 1:1000, 5F7), anti-CyclinA (Santa Cruz Biotechnology, 1:250, sc-596), anti-GAPDH (Santa Cruz 1:5000, D16H11), anti-pS139 H2AX (Cell Signaling 1:300, 9781S), anti-Vinculin (Cell signaling 1:5000, 13901T).

DNAPKi (NU7441, Sigma Aldrich) was dissolved in DMSO and given to cells at a concentration of 2µM 1h before DNA damage induction.

RAD51i (B02, Sigma Aldrich) was dissolved in DMSO and given to cells at a concentration of 27µM l 1h before DNA damage induction.

### Protein Extracts and Immunoblotting

Cells were lysed in IP Buffer (50mM Tris-HCL pH 7.5, 250mM NaCl, 1%NP40, 5mM EDTA, 5mM EGTA) supplemented with 1mM phenyl-methyl-sulfonyl-fluoride, 25mM NaF, 1mM sodium orthovanadate, 25mM β-glycerol-phosphate, 10mg/ml TPCK, 10mg/ml cocktail inhibitor. Alternatively, cells were lysed in RIPA Buffer (50mM Tris-HCL pH 8.0, 150mM NaCl, 1%NP40, 12mM sodium deoxycholate) supplemented with the above described inhibitors.

For immunoblotting 20-50 µg of protein extract were separated by SDS-PAGE, blotted into nitrocellulose membrane and blocked with 5% skimmed-milk or BSA. Immunoblots were developed using ECL (Bio-Rad) or ECL Select (Amersham) using the iBright CL750 imaging system (Thermo-Fisher Scientific).

### Immunofluorescence

Cells were plated on ethanol-washed cover-slips and grown at 37°C and 5% CO_2_ for at least 24 hours. Cells were washed with 1x phosphate buffer saline and fixed in 4% paraformaldehyde 10 minutes at room temperature. Cells were permeabilized with a 0.25% PBS-triton solution for 5 minutes at room temperature. For γH2AX staining cells were additionally permeabilized in cold methanol for 10 minutes at -20°C. Unspecific signal was blocked with 3% bovine serum albumin (Sigma Aldrich) for 1 hour at room temperature. Primary antibodies were incubated 2 hours at 37°C or overnight at 4°C. Secondary antibodies (Alexa Fluor 488, Cy5, Jackson ImmunoResearch) were used 1:300 for 45 minutes at 37°C. Nuclei were stained with Hoechst 33342 for 10 minutes. Images were acquired at a confocal microscope (Zeiss LSM 800) or at a fluorescence microscope (Zeiss). Antibodies used: 53BP1 (Genetex HL275 1:250 – overnight), BRCA1 (Santa Cruz Biotechnology 1:250 - overnight), Cyclin A (Santa Cruz Biotechnology 1:250 - overnight), γH2AX (Cell Signalling 1:300 - 2 hours).

### Neutral Comet Assay

The amount of damaged DNA present in the cells was evaluated by Neutral Comet assay (single cell gel electrophoresis) in non-denaturing conditions. This assay reveals the amount of double-strand breaks present in cells. Briefly, microscope slides were washed in water and then ethanol to remove dust and fatty residues. Slides were then covered twice using Low Melting Point agarose (LMA) previously dissolved at 37°C and left to dry overnight. Cells were plated at a concentration of 4X10^5^ and grown for 24 hours before treatment with 5Gy IR. Pellets were collected at the indicated time-points and stored at -80°C. Subsequently, pellets were suspended in PBS and kept on ice to inhibit DNA repair. Cell suspensions were rapidly mixed with LMA, previously kept at 37 °C and an aliquot was pipetted onto the agarose-covered microscope slides. Cells on the coated slides were lysed for 1 hour at 4°C in the dark (30 mM EDTA, 0,1% SDS). After lysis, slides were washed in TBE 1X buffer (Tris 90 mM; boric acid 90 mM; EDTA 4 mM). Electrophoresis was performed for 20 min in TBE 1X buffer at 0.5 V/cm. Slides were subsequently washed in ddH2O and finally dehydrated in ice cold methanol. Nuclei were stained with GelRed (1:1000 Sigma Aldrich) and visualized with a fluorescence microscope (Zeiss), using a 40X objective. At least 300 comets per cell line were analyzed using Comet-Score software.

### MTS assay

U87-MG and A172 cell lines were pre-treated with inhibitors as indicated above and then treated with 10Gy IR. After radiation cells were plated 1000 per well in 96-well plates and grown for 48h. Cellular viability was measured through Cell titer 96 AQueous One Solution Cell Proliferation Kit (Promega). A yellow solution containing a tetrazolium compound (MTS) and an electron coupling reagent (PES) was added to the cell medium and colour variation was measured after 2 hours at 490 nm using a VICTOR Multilabel plate reader.

### Cytofluorimetric analysis

200.000 cells were plated in petri Dish60 mm and treated after 24hours (DMSO, Olaparib 10μM, IR 10Gy, Olaparib plus IR). Cell death was evaluated 72h after IR by using a CytoFLEX S (Beckman Coulter) instrument. Cells were collected, centrifuged for 5’ at 300g, and double-stained by using Annexin V-APC kit, according to the manufacturer’s instructions (eBioscienceTM Annexin V Apoptosis Detection Kits; Thermo Fisher Scientific) and DAPI. Unstained samples were used as control. Quality control was evaluated using CytoFLEX Daily QC Fluorospheres (Beckman Coulter). FCS files were analyzed using CytExpert version 2.2 software (Beckman Coulter). Dead cells (Annexin V+/DAPI+ cells) were graphed as a percentage to control conditions.

### Traffic light Reporter

The traffic light (Sce target) reporter system was generated in U87-MG cells by lentivirus-mediated transduction and cells were selected using puromycin (2µg/mL). Reporter cells were seeded subconfluently and induced for 72 hours in triplicates with lentivirus vectors carrying partial GFP templates prior readouts. Analysis was performed using the Cytek Aurora flow cytometer and results visualized via FlowJo. Three independent experiments were performed and statistical significance was examined by paired t test (Graphpad Prism).

### *In vivo* experiments

#### *Drosophila* strains and rearing conditions

*Drosophila* stocks were maintained on standard fly food (25 g/L corn flour, 5 g/L lyophilized agar, 50 g/L sugar, 50 g/L fresh yeast, 2.5 mL/L Tegosept [10% in ethanol], and 2.5 mL/L propionic acid) at 25 °C in a 12 h light/dark cycle. All fly strains used come from the Bloomington Drosophila Stock Center (BDSC, Indiana University, Bloomington, IN, USA): *w^1118^* (BDSC #3605) and *dredd^Δ44^* (BDSC #80924). The *dredd^Δ44^/dredd^Δ44^* homozygous flies are viable and fertile.

#### Immunostaining and γ-H2Av foci detection

To induce DNA damage third instar larvae were irradiated with 5 Gy of X rays (X-ray machine Gilardoni MHF200 MD); for immunostaining, larval brains were dissected and fixed in 3,7% formaldehyde at various post-irradiation times 10, 30, 60, and 120 min. Brain preparations were then rinsed several times in PBS 0.1% Triton (PBST), blocked 1h at room temperature and incubated overnight at 4° with mouse anti-H2A.v-P 1:20 in BS (DSHB AB_2618077). The day after rinsed in PBST and incubated for 2h at room temperature Rhodamine conjugated goat anti-mouse 1:50 (Jackson). All slides were mounted in Vectashield medium H-1200 with DAPI to stain DNA. Brain preparations were analyzed on a fluorescence microscope (Zeiss Apotome) and images acquisition performed using the Zen Pro software (Zeiss). Number gH2A.v foci count was performed using Fiji software (National Institute of Mental Health, Bethesda, Maryland, USA).

#### Immunoblotting

To determine the kinetics of H2Av phosphorylation, larvae were irradiated with X rays (5 Gy) and 20 brain samples were collected at different Post IR times (0, 10, 30, 60 and 120 min). Brains were lysed in in sample buffer (Lammeli buffer 2X), fractionated by SDS-PAGE and transferred to nitrocellulose membrane. Primary antibodies were anti-H2Av-P mouse (1:500; DSHB AB_2618077) and anti-vibrator rabbit (1:5000; [35]).

### Statistical Methods

All data were analysed and presented as mean of three independent experiments ± SD, except where differently specified. Significance was determined using Graph-Pad software, performing a two-tailed Student’s t-test, or a two-way ANOVA. P-value < 0.05 was considered significant (*), p-value < 0.01 was considered very significant (**) and p-value < 0.001 was considered extremely significant (***). The functional enrichment analysis was performed by querying EnrichR tool [36].

## ACNOWLEDGEMENTS

We thank all lab members for critical reading of the manuscript and helpful discussion. This work has been supported by research grants from Associazione Italiana per la Ricerca sul Cancro AIRC-IG2016-n.19069, AIRC-IG2021-n.26230, and Italian Ministry of Health, RF-2016-02362022 to D.B.; A.F. phD was supported by MUR, while her PostDoc fellowship was supported by American Italian Cancer Foundation Post-Doctoral Fellowship Award; C.Co. work is been supported by AIRC-IG2021-n.26230; C.D.G. is supported by a MUR fellowship DM 118/2023 (Next-GenerationEU – "Piano Nazionale di ripresa e Resilienza" italiano) to the PhD Program in Cellular and Molecular Biology, Department of Biology, University of Rome Tor Vergata; C.Ci work has been supported by AIRC IG2016-n.19069, AIRC-IG2021-n.26230 and FIRC488 AIRC fellowship for Italy “Filomena Todini”; D.D.B is supported by AIRC IG 2020 - ID. 24315 project.; G.F. work was supported by the cofounding of “Unione Europea – FSE REACT-EU, PON Ricerca e Innovazione 2014-2020 – DM 1062/202” and by Finalizzata Giovani Ricercatori 2021 (grant n.: GR-2021-12372614, CUP: B83C22007560001).

## AUTHOR CONTRIBUTIONS

A.F. performed the experiments on U87-MG and GBM39 cells, data analysis, interpretation and contributed to write the article; C.Co. performed the experiments on A172 and MEF cells, the micronuclei experiments, the chromatin purification experiments, the experiments of synthetic lethality, data analysis, interpretation and contributed to write the article; C.D.G. performed the immunofluorescence analysis of Caspase-8 localization on U87-MG and A172 cells and the cytofluorimetric analyses on U87-MG cells; C.Ci. and D.D.B. evaluated the data and contribute to write the article; G.F. and P.P. produced the bioinformatics data; M.G. irradiated cells; M.M. and L.C, conceived and performed the Drosophila experiments and analysis; T.Y. and R.S. performed the experiments with the traffic light reporter system; D.B. designed the experiments, evaluated the data and wrote the paper.

## DATA AVAILABILITY

All data in the manuscript are available in the Supplementary data, and raw data are available upon request. The transcriptomic analyses reported in this manuscript are based on previously published RNA-seq experiments (Contadini et al., 2023, Cell Death Differ) and the raw RNA-seq data can be accessed at GEO, Go to https://www.ncbi.nlm.nih.gov/geo/query/acc.cgi?acc=GSE193495 Enter token sjojmgoefvgflqd into the box. The proteomic analyses reported in this manuscript are based on an article recently accepted for publication (Cirotti et al., 2025, CDD-24-2998RR) and the raw data are available via ProteomeXchange with identifier PXD060920 (Token: s5h584Mp3CfY).

